# A Modular Microarray Imaging System for Highly Specific COVID-19 Antibody Testing

**DOI:** 10.1101/2020.05.22.111518

**Authors:** Per Niklas Hedde, Timothy J. Abram, Aarti Jain, Rie Nakajima, Rafael Ramiro de Assis, Trevor Pearce, Algis Jasinskas, Melody N. Toosky, Saahir Khan, Philip L. Felgner, Enrico Gratton, Weian Zhao

## Abstract

To detect the presence of antibodies in blood against SARS-CoV-2 in a highly sensitive and specific manner, here we describe a robust, inexpensive ($200), 3D-printable portable imaging platform (TinyArray imager) that can be deployed immediately in areas with minimal infrastructure to read coronavirus antigen microarrays (CoVAMs) that contain a panel of antigens from SARS-CoV-2, SARS-1, MERS, and other respiratory viruses. Application includes basic laboratories and makeshift field clinics where a few drops of blood from a finger prick could be rapidly tested in parallel for the presence of antibodies to SARS-CoV-2 with a test turnaround time of only 2-4 h. To evaluate our imaging device, we probed and imaged coronavirus microarrays with COVID-19-positive and negative sera and achieved a performance on par with a commercial microarray reader 100x more expensive than our imaging device. This work will enable large scale serosurveillance, which can play an important role in the months and years to come to implement efficient containment and mitigation measures, as well as help develop therapeutics and vaccines to treat and prevent the spread of COVID-19.

## Introduction

To date, the 2020 COVID-19 pandemic has claimed hundreds of thousands of lives, with many more to come, shattered health care and social systems and crippled the economy on an unprecedented global scale^1^. Long incubation periods in combination with transmission through pre-symptomatic and asymptomatic carriers, exacerbated by the highly contagious nature of SARS-CoV-2, have rendered prevention of community spread very difficult^2–11^. As an essential step towards recovery, to implement efficient containment measures, and to help develop therapeutics and vaccines, we must implement broad testing, for the virus and for antibodies against the virus.

To better understand the humoral response to viral exposure, model the spread of COVID-19, and help orchestrate local public health containment measures, we recently constructed a novel serology test, Coronavirus Antigen Microarray (CoVAM)^12,13^. CoVAM can currently measure antibody levels in serum samples against 67 antigens from 23 strains of 10 viruses known to cause respiratory tract infections including SARS-CoV-2. New antigens can be included as the virus evolves. Probing this large number of antigens simultaneously in a single test allows for much higher specificity, sensitivity, and information density than conventional antibody tests such as lateral flow assays (LIFAs). LIFAs are susceptible to false positive results, especially for COVID-19 and current LIFA test performance has been reported inadequate for most individual patient applications^14^. Testing for reactivity against only one or two antigens is not always reliable as cross-reactivity can occur. The CoVAM test can tease out this cross-reactivity by taking a simultaneous snapshot of the relative serum reactivity against multiple, cross-species viral antigens. In addition, each array contains four replicates of the same set of antigens to vastly improve statistical power. This way, CoVAM can easily discriminate SARS-CoV-2 from SARS, MERS and other common coronaviruses^12,13^. Furthermore, the highly specific CoVAM array is specifically designed for low-cost, high-throughput serological studies on the scale of >100,000 samples, which will be critical as the virus is spreading to low-income countries with large populations.

While microarrays could be printed and distributed on a large scale, reading the slides by fluorescence imaging currently requires expensive ($10,000 – $100,000) machines which many clinical laboratories currently do not possess and are especially difficult to move to makeshift testing sites including field clinics. Sending the probed slides back to designated imaging centers is expensive and time consuming, therefore unsuitable for the required large-scale testing. In the upcoming months and years, serosurveillance technology to mitigate the continuing spread of COVID-19 and other viral pathogens must be capable of repeated testing of a large global population. To make this possible, a robust, inexpensive, portable imaging platform that can be deployed immediately in any basic laboratory to read coronavirus antigen microarrays is required. This will be especially valuable in countries with otherwise highly vulnerable populations due to restricted access to tests and lack of a suitable health care infrastructure. To address this issue, we have developed a robust, inexpensive ($100 – $300), and portable imaging platform, the TinyArray imager, that can be deployed immediately in any basic laboratory. Our TinyArray imager uses a 3D printable design in which widely available components were used to excite fluorescence of labeled secondary antibodies that can be detected with an inexpensive 5-megapixel camera module with sufficient spatial resolution and sensitivity to reliably read microarrays. In this work, we show that, with patient samples, this imaging platform can match the results obtained with a 100x more expensive commercial imager; linear regressions of microarray fluorescence intensities consistently showed R-squared values >0.85 between imaging systems, similar to linear regressions between image replicates acquired on the same device. By bringing low cost, high throughput, highly specific serological testing to the public, our platform could be highly valuable for COVID-19 serosurveillance.

## Results

### Imager Design

Various low-cost microscopy platforms using portable devices such as cell phone cameras have been recently developed for various applications. In mobile devices, different illumination strategies have been reported including on-axis epi-illumination^15,16^, off-axis inclined illumination^17^, butt-coupling^18^, and total internal reflection^19^. In order to avoid out-of-focus background with these illumination schemes, either the sample is compressed to a thickness of ~10 μm by mounting it between two glass slides^20^ or physical properties of the sample such as plasmonic enhancement due to the presence of a metal surface^17^ or total internal reflection due to the presence of a refractive index change are exploited^19^. Yet, to reliably read microarrays, a suitable device also needs to provide a large (5-75 mm), uniform, undistorted field of view with high spatial resolution (~10 μm), and the sensor response across the field of view must be calibrated to ensure a linear response. Also, probing of multiple antibody isotypes such as IgA, IgM, and IgG in a single test requires several spectral windows for illumination and detection of secondary antibodies labeled with different dyes; the light sources must be arranged to ensure consistent, homogeneous illumination and detection. Finally, the imager should be linked to a data bank or computational facility to safely store and analyze patient data. To test a broad population in an inexpensive, high throughput manner, the microarray reader should combine all these traits while being portable, of low cost, and easy to use by non-experts with minimal training. The device should also feature low power consumption to allow battery powered operation and be possible to manufacture on a large scale in a simple fashion. Our TinyArray imager combines all these traits to read microarrays on a large scale (**Fig. 1a**).

**Figure 1.**
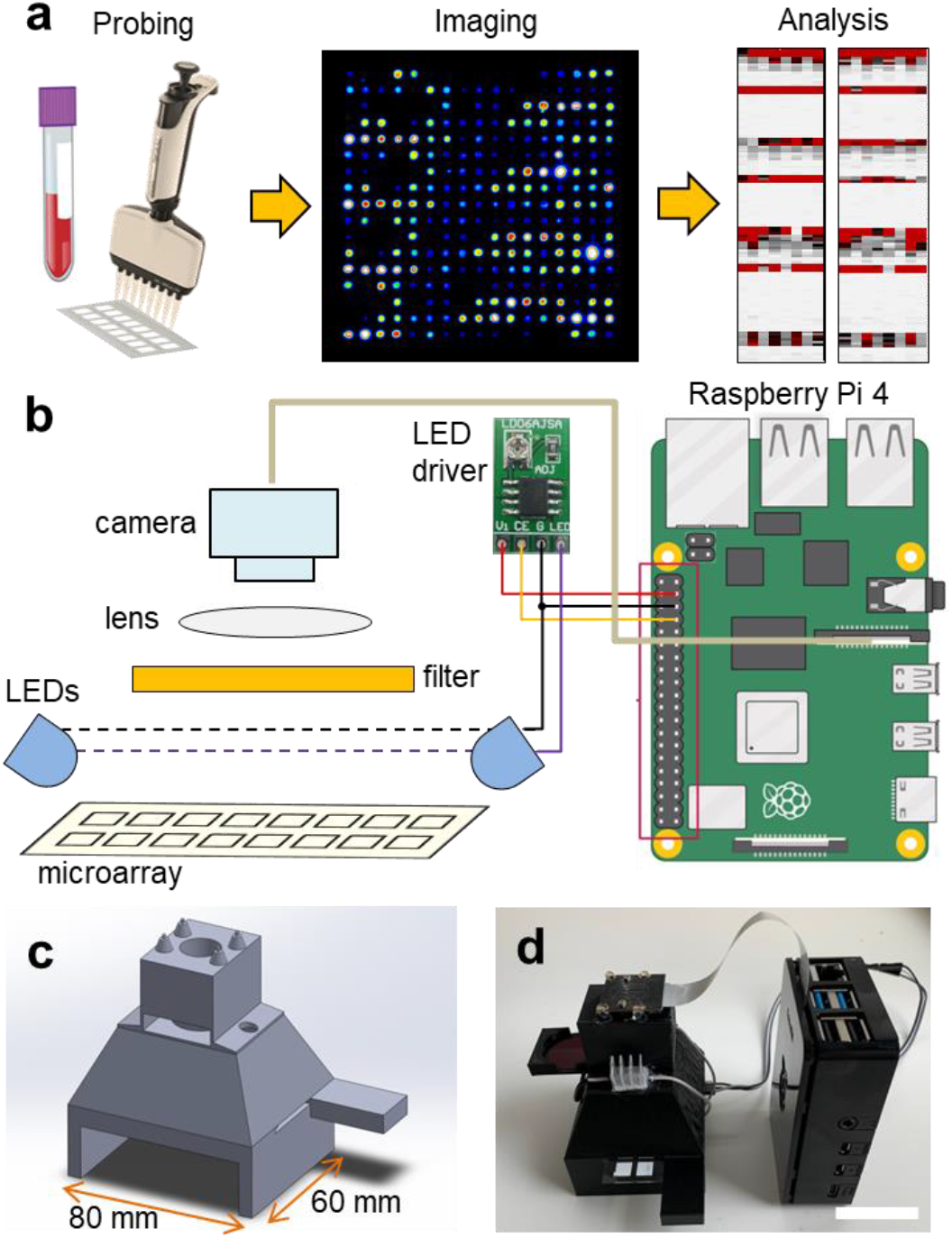
TinyArray imager design. (**a**) Workflow: After probing of the antigen microarrays, images are taken where the fluorescence intensities corresponding to the relative antibody concentration are quantified (**b**) The microarray was LED-illuminated (470 nm) from the top and imaged through long pass and band pass filters with an OmniVision OV5647 sensor module. Illumination was controlled and images were acquired with a single board computer (Raspberry Pi 4). (**c**) CAD design of the microarray imager. (**d**) 3D printed and assembled prototype together with a Raspberry Pi 4 single board computer interfacing the camera and 75 mm × 25 mm × 1 mm microarray slide inserted into the device. Scale bars, 30 mm.

We focused on imaging microscope slides of 25 mm × 75 mm × 1 mm as these are the most common for microarray printing. Other formats could be imaged by adjusting the slide holder size or by using an adapter. As the best compromise between ease-of-use and hardware requirements, the slide is imaged in two steps using a slot-in design.

This way, the slide can be positioned precisely into the camera field of view by inserting it into the reader until it reaches a stop. This process is repeated by inserting the slide from the other end. To image the entire slide as a whole, the resolution of the camera chip and size of optics would need to be very large, while capturing more than two smaller fields of views would require a more complicated slide positioning mechanism. Microarray images are acquired from the top with a 5-megapixel OmniVision OV5647 sensor that is widely available as a camera module for popular single board computers such as the Raspberry Pi (**Fig. 1b**). Placed immediately before the camera lens (3.6 mm focal length), long pass (LP570, LP660) and band pass filters (BG40) were used to spectrally select for the fluorescence emission of the employed fluorophores. Fluorescence excitation was realized from the top with two LEDs (470 nm) along the long axis of the microarray slide next to the camera at a slight angle to ensure homogeneous illumination of the field of view. All components including illumination, detection optics, and the sample holder are integrated in a CAD/CAM model that can be manufactured on a large scale at low cost, for example, by 3D printing or injection molding (**Fig. 1c**). A picture of the 3D printed and assembled microarray imager prototype is shown in **Fig. 1d**.

### Evaluation with Microarrays of Serial Dilutions of Quantum Dot Probes

To evaluate the performance of the imaging device, we printed microarrays with serial dilutions of QD655-Streptavidin, sample fluorescence images are shown in **Fig. 2a,b**. For testing, 16-pad microarray slides (Oncyte Avid, Grace Bio-Labs) with pad dimensions 7 mm × 7 mm were used, and microarray dots in a configuration of 18 × 18 spots per pad (150 μm diameter, 280 μm spacing) were printed with serial dilutions of QD655-Streptavidin across a concentration range of three orders of magnitude (**Fig. 2c**). For evaluation, images of these test slides were acquired with a camera-based system (ArrayCAM 400-S, Grace Biolabs) and our TinyArray imaging module. For illumination, two 470-nm LEDs were used with a battery-powered driver circuit (LD06AJSA, Amazon). Fluorescence was detected through a 570-nm long pass filter (Schott OG570, Thorlabs). Our data show that the 5-megapixel sensor employed has enough spatial resolution and sensitivity to reliably read microarrays. Images were acquired and processed using the Python programming language and related libraries and packages (available at https://www.python.org/). The camera exposure time was adjusted to optimize the dynamic range to the fluorescence signal and to ensure a linear response. Before microarray imaging, reference images of a well-defined 7 × 14 square checkerboard pattern were taken and the camera lens aberrations were measured and corrected using the Open Computer Vision (OpenCV) package (available at https://opencv.org/). Background was removed by subtraction of a median-filtered (15 pixel radius) version of the same image to eliminate spatial variations of the signal offset. Fluorescence signal in the individual dots is not affected by this as they occupy areas less than 15 pixels across. Image data was uploaded to a workstation through the single board computer’s integrated WiFi connection where it was quantified with Mapix (Innopsys), as shown in **Fig. 2d,e**. From the images and the cluster analysis, it can be seen that the data obtained with the TinyArray imager is comparable to the ArrayCam 400-S. To further quantify, we evaluated the TinyArray imager with microarrays that had been probed with serum samples and developed for antibody signals using anti-human IgG and anti-human IgA secondary antibodies that were conjugated to QD800 and QD655, an exemplary raw fluorescence image of 2 × 2 microarray pads is shown in **Fig. 2f**, the corresponding background-subtracted image is show in in **Fig. 2g**. In comparison with data from an ArrayCAM 400-S commercial imager, we obtained R^2^ >0.85 through linear regression (**Fig. 2f**). This is similar to differences between image repeats of the same samples taken by the ArrayCam and confirms that it is possible to use our simple design instead of expensive, bulky, lab-based laser scanners.

**Figure 2.**
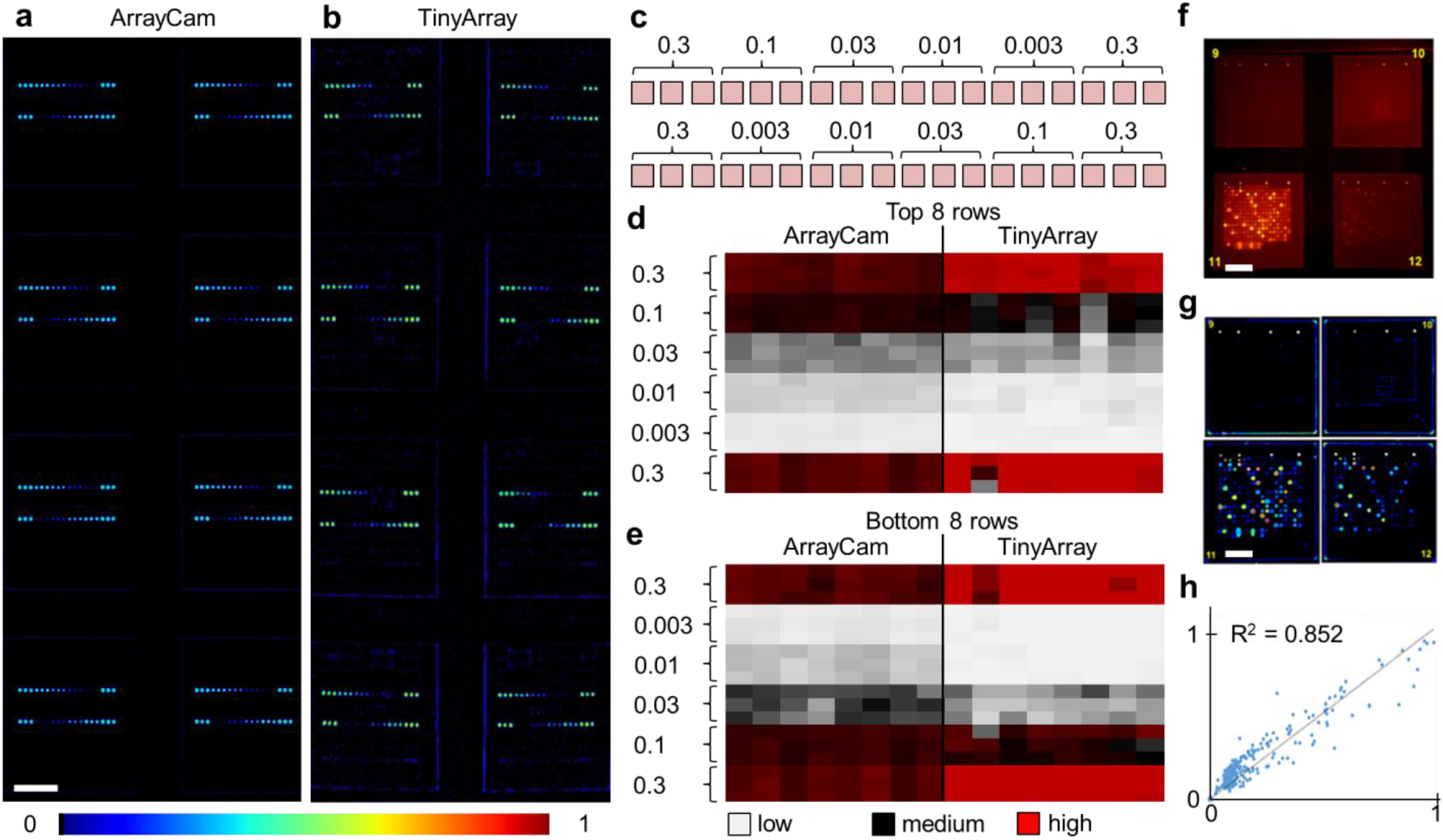
Fluorescence images and quantification of printed microarrays with 280-μm spaced, 150 μm-diameter dots of labeled with quantum dot probes. (**a,b**) Fluorescence images of 2 × 4 microarray pads (7 mm × 7 mm each) taken with the commercial ArrayCam 400-S and the TinyArray imager, respectively. Fluorescence intensity is represented on a pseudo rainbow color scale (**c**) Layout and relative concentrations of the serial dilutions of QD655-Streptavidin microarray dots imaged in panels (a,b). (**d,e**) Quantitative analysis of the background-subtracted median intensities in the top (panel d) and bottom QD655-Streptavidin dot rows (panel e) of the images taken with the ArrayCam (left column) and the TinyArray imager (right column) as shown in panels (a,b). (**f**) Raw image of four microarray pads probed with human serum samples and developed with secondary antibodies conjugated to QD655. (**g**) Background-subtracted microarray image in pseudo rainbow color scale. (**h**) Graph and linear regression (R^2^ >0.85) of the background-subtracted median fluorescence intensities of the same microarray sample as obtained with the TinyArray imager prototype and the ArrayCam 400-S. Scale bars, 2 mm.

### Evaluation with Coronavirus Microarrays

We previously used CoVAM to detect IgG and IgA antibodies against a panel of antigens including coronavirus spike protein (S) as separated receptor binding (RBD), S1, and S2 domains or whole protein (S1+S2) and nucleocapsid protein (NP) from multiple coronaviruses including SARS-CoV-2, SARS-CoV, MERS-CoV, and common cold coronaviruses (HKU1, OC43, NL63, 229E)^12,13^. In Assis et al., high-performing antigens for IgG and IgA detection, defined by Receiver Operating Characteristic Area Under Curve (ROC AUC) >0.95, were found to discriminate between the positive group and negative groups with high significance including all SARS-CoV-2 antigens and MERS-CoV S2 for IgG and SARS-CoV-2 S2 and S1+S2 for IgA. As the microarray reader was specifically designed to enable broad access to microarray-based SARS-CoV-2 antibody testing, we repeated probing of a cohort of samples and imaged the corresponding microarrays with our TinyArray imager prototype and the ArrayCAM 400-S commercial imager. Coronavirus microarray slides were developed with secondary antibodies against human IgG (Qdot 800 conjugated anti-human IgG, CAT#110610, Grace BioLabs) labeled with quantum dot fluorescent probes as previously described^12,13^. Example images of microarrays probed with positive sera are shown in **Fig. 3a** (ArrayCAM 400-S) and **Fig. 3b** (TinyArray imager). The background-subtracted median fluorescence intensities were quantified for each spot. The normalized intensities measured with the TinyArray imager were plotted against the values obtained with the ArrayCAM 400-S in **Fig. 3c**; linear regression resulted in R^2^ values of 0.82-0.92. CoVAM heat maps of the 5 SARS-CoV-2-positive (9 arrays including 4 duplicates) and 10 negative control samples were generated from the Array Cam 400-S and TinyArray imager data as shown in **Fig. 3d**. Antibodies in positive sera binding to the seven SARS-CoV-2 antigens were further quantified in **Fig. 3e** for the Array Cam data and **Fig. 3f** for the TinyArray imager data.

**Figure 3.**
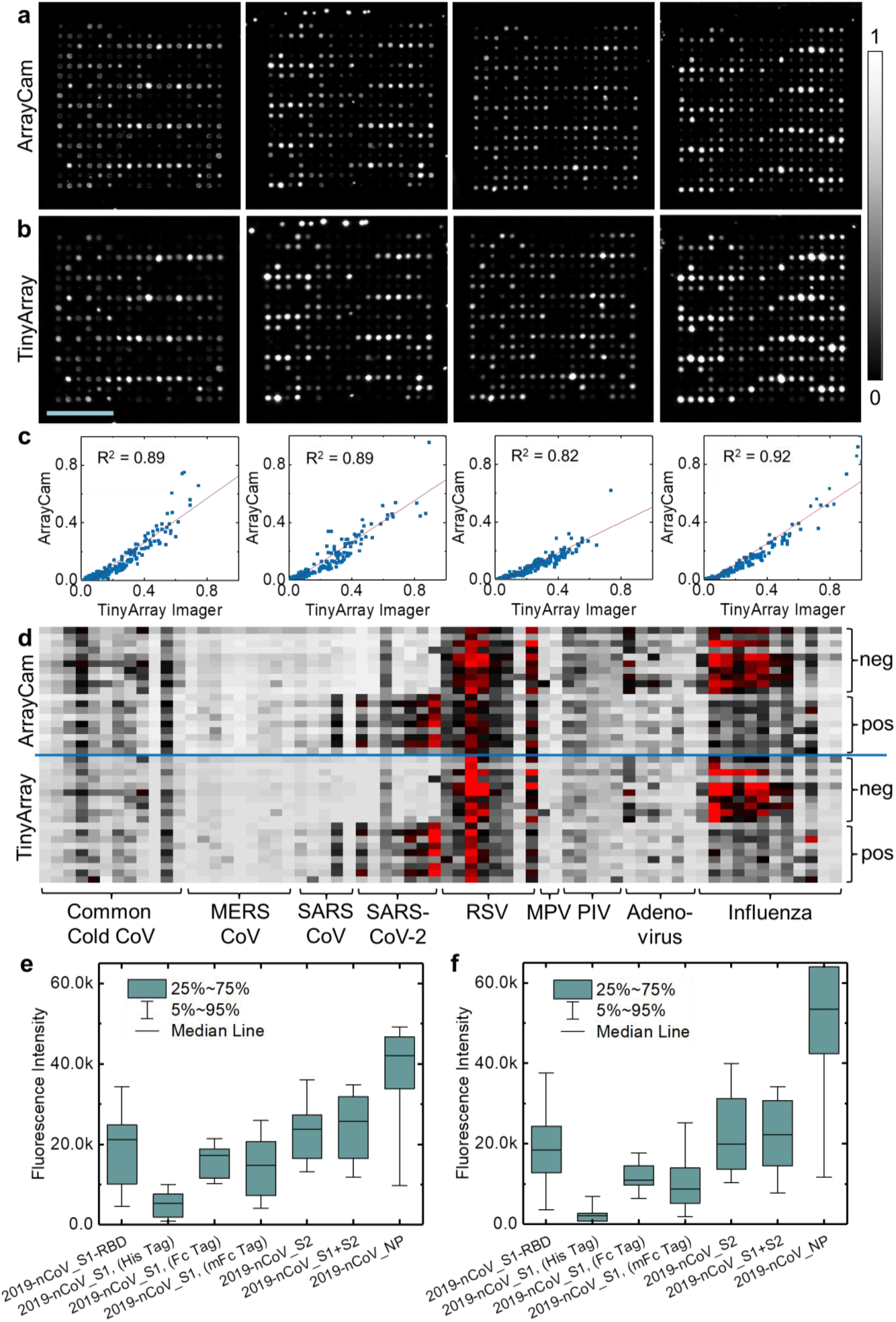
Fluorescence images and data analysis of CoVAM probed with positive sera. (**a**) Four exemplary fluorescence images acquired with the Array Cam 400-S. (**b**) Corresponding TinyArray images. (**c**) Background-subtracted median fluorescence intensities obtained for each microarray spot with the Array Cam 400-S and the TinyArray imager that were normalized and plotted against each other; linear regressions were performed and R^2^ values were calculated. (**d**) Heat maps of 9 SARS-CoV-2-positive and 10 negative control samples generated from the Array Cam 400-S (top row) and TinyArray imager data (bottom row). Gray/black/red colors indicate low/medium/high antibody prevalence. (**e,f**) Statistical analysis of the seven SARS-CoV-2 antigens in the CoVAM for positive sera for the ArrayCam and the TinyArray imager data. Scale bar, 2 mm.

## Discussion

It is well accepted that official infection numbers of COVID-19 are widely underestimated. This is due to a shortage of tests, limiting testing to people with symptoms and the time-sensitive nature of RT-PCR as it depends on the presence of viruses and/or viral genetic material in respiratory tract mucosa. Broad availability of highly specific, high-throughput, inexpensive serological testing can help manage COVID-19 over the coming months and years as it will be able to determine the true density of exposed, seropositive people to enable containment and mitigation measures to avoid formation of new COVID-19 hot spots. Massive serological testing could also help devise strategies to restart the economy in a controlled way, minimizing the risk of further waves of infections and COVID-19 related deaths. Furthermore, our assay can identify potential blood donors who have recovered from COVID-19, thereby enabling plasma transfusion to treat COVID-19 patients via passive immunization^21^. Lastly, our serology testing can reveal information about the global host immune response to SARS-CoV-2 and provide insights to guide therapeutic and vaccine research and development.

In order to get a comprehensive view of the serological status of a large population as well as better predict disease distribution and outcome to help develop more efficient control measures, microarrays are an invaluable tool. CoVAM is specifically designed for high-throughput serological studies on the scale of >100,000 samples with a minimal number of reagents, which will be critical to enable massive, repeated testing of large populations. For this purpose, testing can be highly parallelized using multi-pad slides or well plates where 16 or 96 patient samples can be probed simultaneously per plate. Imaging with the TinyArray imager only takes tens of seconds such that many slides/plates can be imaged quickly after parallel incubation to scale up throughput. As a robust and inexpensive ($200) imaging platform, we can deploy this powerful technology anywhere in the world to fight COVID-19. After imaging, microarray data could be uploaded for cloud-based analysis using a smartphone. This capability will be especially important in the upcoming months as the disease is spreading to countries with minimal health care infrastructure and high population densities.

## Materials and Methods

### TinyArray Imager Design

The sample slides were illuminated with two 3-W LEDs (2 × 365 nm or 2 × 470 nm) from the top; LEDs were driven by a DC-DC driver circuit to adjust the current (max 300 mA). LED illumination was switched with the Raspberry Pi GPIO through a MOSFET switch. All electronics were powered by a 5 V, 3.5 A power supply. Images of microarray slides were acquired with a 5-megapixel OmniVision OV5647 sensor without IR filter coupled to a lens of 3.6 mm focal length adjusted to a field of view of 35 × 26 mm. Fluorescence was detected through a combination of a 570-nm long pass filter (Schott OG570, Edmund Optics, Barrington, NJ) with a 335 – 610 nm band pass filter (Schott BG40, Thorlabs, Newton, NJ) to select for QD585 emission and 630-nm/660-nm long pass filters (Schott RG630, Thorlabs/Hoya R-66, Edmund Optics) to select for QD655/QD800 emission. The camera was controlled by a Raspberry Pi 4 single board computer running Raspian 4.19 with software written in Python 3.7. A 3D CAD/CAM model to house the components and hold the microarrays slides was created in Solidworks 2019/2020 (Dassault Systemes, Waltham, MA). After 3D printing of the model (Ender-3 pro, Creality3D, Hong Kong), all relevant optical components were inserted and attached. The microarray slides used consisted of 2 × 8 pads of 7 mm × 7 mm each (Oncyte Avid, Grace Bio-Labs, Bend, OR). In this configuration, 2 × 4 pads were imaged simultaneously with ample spatial resolution (13.5 μm sample pixel size) to resolve the individual dots (150 μm diameter, 280 μm spacing) of the 18 × 18 dot microarrays. Hence, the whole 2 × 8 array could be read by taking only two images. The microarray slides could be inserted directly into the device in a dedicated slot that maintains the correct distance/position to the illumination and detectors (here, 40 mm). After slide insertion into the slot in either direction, bump stops at the end ensure the correct position (pads 1-8 and pads 9-16, respectively) of the slide to reliably image all pads and to make sure that the individual pads of the slide were indexed correctly.

### Image Acquisition and Data Transfer

Software deployed on the microarray imaging device ensured acquisition of reproducible, quantifiable images of antigen arrays. In the prototype, we used the Python programming language and the following libraries: picamera, numpy, Image, scipy. Python and packages are freely available at https://www.python.org/. Images were transferred through the Raspberry Pi’s internal WiFi connection. Camera exposure times (1s – 5 s) and sensitivity settings (ISO 100 – ISO 800) were adjusted to optimize the dynamic range to the fluorescence signal and ensure a linear response. To reliably quantify microarrays, it is important that the individual spots appear in a defined grid at constant distances. Therefore, spherical aberrations of the imaging lens were characterized by taking a reference image of a checkerboard pattern (7 × 14 squares) and corrected in the microarray images using the Open Computer Vision (OpenCV) package for Python (https://opencv.org/). Background was subtracted to eliminate spatial variations of the signal offset. For this purpose, a 15-pixel median filter was applied to the image data and the result subtracted from the raw image.

### Microarray Image Data Quantification

To enable quantification of the dye concentration indicative of the antibody concentration in all spots of the array, the intensity must be determined in each spot. For this purpose, microarray slides were printed with a distinct pattern of 18 × 18 dots (150 μm diameter, 280 μm spacing). The slides were imaged using both the TinyArray imager and the commercial ArrayCam system (Grace Bio-Labs, Bend, OR). As the size and distances of the spots were known, the background-subtracted median spot fluorescence intensity was measured and quantified using Mapix (Innopsys) as previously described^22–24^. Mean fluorescence intensity (MFI) of the four replicate spots for each antigen was utilized for antigen heat map generation and data analysis.

### Coronavirus Antigen Microarray Probing

Coronavirus microarrays (CoVAMs) were probed as previously described^12,13^. CoVAM included 67 antigens against 23 respiratory virus strains including SARS-CoV-2, provided by Sino Biological Inc. (Wayne, PA). Four replicates of antigen patterns were printed in a 18 × 18 dot arrangement onto each 7 mm × 7 mm pad of the 2 × 8 pad nitrocellulose-coated microarrays slides (Grace Bio-Labs, Bend, OR) with an OmniGrid 100 microarray printer (GeneMachines). Before printing, lyophilized antigens were reconstituted to a concentration of 0.1 mg/mL in printing buffer containing protein stabilizers. Each pad was probed with human sera, imaged, and analyzed as previously described^22–24^. Briefly, slides were probed with 1:100 dilutions of human sera with 1× GVS Fast Blocking Buffer (Fischer Scientific) and incubated for 2 h at room temperature. The microarrays were then washed 3x with T-TBS buffer (20 mM Tris-HCl, 150 mM NaCl, 0.05% Tween-20, adjusted to pH 7.5 and filtered), labeled with secondary antibodies to human IgA and IgG conjugated to quantum dot (QD) fluorophores for 1 h at room temperature, followed by a second 3× wash with T-TBS before drying.

### Specimen Collection

Convalescent sera from confirmed recovered coronavirus cases were provided by Ortho Clinical Diagnostics, which is using these specimens to validate a clinical diagnostic test. Evaluation of these samples on the coronavirus antigen array are published^12^. The pre-COVID-19 naïve blood sera were collected in the context of a larger study to identifying acute respiratory infection (ARI) cases in a college resident community in the Eastern United States monitored using questionnaires and RT-qPCR^25^. From among de-identified blood specimens for which future research use authorization was obtained, five specimens that showed high IgG reactivity against human coronaviruses in the larger study were chosen for validation of the coronavirus antigen microarray^13^.

## Data Availability

Upon request, we will make the data available to other researchers. The TinyArray imager CAD file and python program code will be made available for non-commercial purposes upon request.

## Acknowledgements

We thank Byron Shen from Velox Biosystems for help with providing materials. This work was supported by National Institutes of Health grants P41 GM103540, R01 AI117061, and a UC Irvine CRAFT-COVID grant.

## Author Contributions

P.N.H., E.G., W.Z., and P.L.F conceived the project. A.Jain, R.N., R.R.A., S.K., M.N.T. and A.Jasinskas supplied, prepared and probed samples and analyzed data. P.N.H designed, constructed and tested the TinyArray imager and wrote software. T.J.A. acquired image data. T.P. helped design the CAD model. P.L.F., W.Z., and E.G. supervised the project and supplied materials. P.N.H. wrote the manuscript and coordinated the project.

## Author Disclosures

The coronavirus antigen microarray is intellectual property of the Regents of the University of California that is licensed for commercialization to Nanommune Inc. (Irvine, CA), a private company for which Philip L. Felgner is the largest shareholder and several co-authors (de Assis, Jain, Nakajima, Jasinskas, and Khan) also own shares. Nanommune Inc. has a business partnership with Sino Biological Inc. (Beijing, China) which expressed and purified the antigens used in this study. Weian Zhao is the founder of Velox Biosystems Inc. (Irvine, CA) that develops diagnostics including for COVID-19.

